# Distinct neural markers for intentional and unintentional task unrelated thought

**DOI:** 10.1101/705061

**Authors:** Adrien Martel, Mahnaz Arvaneh, Ian Robertson, Jonathan Smallwood, Paul Dockree

## Abstract

Studies suggest that generating thought unrelated to the task in hand is accompanied by a reduction of attention to external task-relevant information. This observations led contemporary theory to suggest multiple component processes contribute to patterns of ongoing thought. The present study used EEG to seek support for these component-process accounts by examining the neural correlates of deliberate and spontaneous task unrelated thought. EEG activity was compared prior to reports of ongoing thought during a test of sustained attention. Event-related potentials, such as the P3, were attenuated during off-task states, regardless of whether they were intentional or not. In contrast, increased alpha power and cortical phase-locking were linked to deliberate off-task thoughts, while reductions of evoked sensory response were prevalent in spontaneous off-task episodes. These data suggest off-task thought shares common and distinct neural features that are differentiated through their relationship to intention, supporting component process accounts of ongoing thought.

## Introduction

Ongoing thought is not always directed to events taking place in the external environment or the task being performed, but is often devoted to information generated from internal representations, such as episodic or semantic memories^1–4^ These off-task experiences are common in the lab^5–8^ and daily life^9–12^, and can vary in their content^13–16^ and more abstract features such as metacognitive awareness^17–19^ and their links with intention^20–23^. Recent views from both psychology^24^ and neuroscience^25^ have argued that off-task experiences can be understood as a family of states with overlapping experiential neural and cognitive features.

One approach to explaining the apparent heterogeneity of the off-task state assumes that different types of experience emerge from variation in the contribution of distinct underlying neurocognitive processes^4^ Evidence for this *component process* account of ongoing thought come from studies showing both common and distinct neuro-cognitive profiles for different aspects of the off-tasks state. For example, studies have found that off-task thinking shows reduced attention to external task relevant information as measured by event related potentials, regardless of the level of awareness of ongoing experience^26^ and relies on a similar structural and functional architecture regardless of whether focused on the past or the future^2,27^. In contrast, features of off-task thought, such as its level of intentionality, may have their roots in distinct neurocognitive features. For example, trait variation in spontaneous off-task thought is linked to symptoms of attention deficit disorder^28^, is more prevalent in demanding tasks^29^ while individuals whose off-task thought are more deliberate show greater cortical thickness in areas of cortex thought to be linked to executive control^30^.

There are at least two processes that contemporary accounts of off-task thought assume to be important during off-task states^31^. Since off-task thoughts are related to information that is absent from the external environment, these experiences are assumed to rely on the ability to decouple attention from events in the external environment. This enables the ability to focus on information not present in the immediate environment and is thought to be necessary for off-task thoughts because of the limited capacity of attentional processing^32^. In contrast, the process of executive control is important in constraining the focus of attention to information relevant to a particular train of thought, and is thought to support both task relevant and off-task patterns of cognition depending on either the demands of the task^33,34^, or, the motivation of the individual^30^ see also^35^. Unlike perceptual decoupling, the process of executive control is thought to contribute to off-task states when they are engaged in a deliberate manner.

In this past decade, the electrophysiological dynamics underlying periods of on- and off-task thought has been the object of numerous studies linking perceptual decoupling with characteristic changes in the time and frequency domain of the EEG. One consistent finding is that cortical processing of perceptual input, as measured by electrophysiological indices, is transiently dampened during off-task mental activity^26,36–39^. An effect that seems to support the idea that the ability to engage in off-task thought hinges on the capacity to decouple attention from the external environment in favor of internal thought processes. Perceptual decoupling has been suggested to be the result of an insulation mechanism that shields the current train of thought from the potential disruption of processing incoming sensory information^4,31,32^. Accordingly, the reduction in both the perceptual- and cognitive-level processing of sensory input during task-unrelated thoughts has been proposed as a likely cause behind the disruption in performance observed across various attentional tasks^5,40,41^. In terms of event-related potentials (ERPs), a reduction in cortical processing was observed across sensory modalities during both behavioral lapses and periods of reported mind-wandering. Early visual components such as the P1, thought to index the suppression of irrelevant stimuli^42^, was observed to be asymmetrically reduced in the right hemisphere^43^ while performing a reading task^39^ contralaterally to peripheral stimuli and bilaterally^36^ during a sustained attention to response task (SART).

Interestingly, electrophysiological evidence across these studies seem to suggest that early perceptual components are only sensitive to modulation with respect to primary task stimuli^36,39,43^. ERPs of secondary task or control stimuli which are outside of the immediate spotlight of attention such as SSVEP^44^ or auditory probes^37,45^ were not modulated by attentional states. Given that early sensory components are known to be modulated top-down by attention^46^, these findings seem to suggest that the mechanisms of perceptual decoupling are engaged in a top-down manner when the sensory input represents a potential threat to the integrity of the current train of thought. J. Smallwood^31^ proposed that the lateral prefrontal cortex (LPFC), a region of the cortex suspected to be involved in the management of endogenous and exogenous attentional states^47–49^ as well as resolving competing stimulus inputs^50,51^, might play a crucial role in regulating sensory input and fluctuations in attentional states. Incidentally, direct current stimulation of the LPFC has been found to have a modulating effect on the propensity for mind-wandering^52–54^.

Returning to the hypothesis that task-unrelated thoughts eventuate a general disengagement of attentional systems from the external environment, the magnitude of components related to higher order perceptual, such as the P2, have recently been observed to be dampened by mind-wandering during semantic encoding^55^. And similarly, the P3, a component thought to reflect the general level of cognitive processing or allocation of attentional resources applied to task stimuli^56,57^, has been shown to be reduced during mind-wandering. Significant reductions in P3 ERP amplitudes during off-task states were observed during a SART^26^, a visual oddball task^38^, the evaluation of complex affective visual stimuli^58^ and during a driving task^37^. Moreover, reduction of the P3 amplitude, which can be considered an index of perceptual decoupling or the level of attention directed towards the task at hand^59^, was observed foreshadowing behavioral lapses in a temporal expectancy task^44^ and during covert sustained attention tasks^60,61^.

As for oscillatory activity, several studies observed increases in alpha activity prior and during attentional lapses^44,59,61–63^ or during periods of reported off-task thought^36^. Although alpha (~8-14 Hz) is the most prevalent rhythm in the human cortex it is suggested to be related to attentional processes^64^, an inverse correlate of cortical excitability, i.e. a gating mechanism actively inhibiting competing or distracting inputs^65,66^. Generally speaking, increased alpha can be seen as an index of the degree to which sensory input is being inhibited in favor of more task-relevant information. In contrast, theta and more specifically frontomedial theta (FMT) was found to be a strong correlate of cognitive effort or mental fatigue in sustained attention^67,68^ and to be upregulated when performing tasks with high demands on externally directed attention^69,70^ or during flow, as state commonly associated with a deep engagement with the task at hand and subjective loss of reflective self-consciousness^71^. With respect to mind-wandering theta activity was found to be inversely correlated with blood-oxygen-level dependent activity in the default-mode network^72^ (DMN), a region extensively linked with internally directed attention^73,74^. At the core of this distributed large-scale network^75^, the posterior cingulate cortex and its connectivity with regions supporting cognitive control^76–78^ and regions supporting episodic knowledge^79–81^ are thought to orchestrate the ongoings of spontaneous thought^2^. Indeed, a recent study in a clinical population showed a correlation between the integrity of this connectivity and mind-wandering capacity^82^. Further, a diffuse increase in theta activity was recorded during mind-wandering as compared to a breath focus task^45^. In the context of neural oscillatory activity, the phase angles reflect the congruence in the timing of stimulus-evoked responses which is quantified with inter-trial phase coherence (ITPC^83^). Theta ITPC, interpreted as a measure of attentional stability^84^, was found to be decreased in the parietal cortex for the P1 time-window during mind-wandering^36^. Importantly, the amplitude of the P1 component was correlated with theta ITPC suggesting a potential role for phase coherence in perceptual decoupling and possibly form an explanation for the consistency of early sensory responses to non-primary task stimuli during off-task thought. In accordance with these findings, off-task states were associated with an increase in power and ITPC for alpha, theta and gamma across regions of the DMN^85^. Taken together, these studies converge on the notion that task-unrelated mental activity attenuates sensory input that could otherwise disrupt its unfoldment. One of the questions remaining is whether the electrophysiological signature of different component processes vary when off-task states are engaged in deliberately or arise spontaneously.

Our study set out to explore the common and distinct neurocognitive processes that underlie patterns of intentional and unintentional off-task states, deliberate (dTUT) and spontaneous task-unrelated thought (sTUT) respectively. In our experiment, we used SART^86^ and used “experience sampling” (ES) to fluctuations in the focus of attention over time^19^. For our experiment, we employed the fixed version of the SART which displays the digits sequentially rather than randomly. The resulting predictability of the stimuli sequence, and thus of the GO/NOGO responses required, should render the task more monotonous, resulting in more lapses of attention^44,87,88^ but also in an increased number of dTUT episodes^23^. Electroencephalography (EEG) was used to record ongoing brain activity during this period. Using these data, we explored the neural processes that take place during periods of off-task thought, as well as the neural patterns that are specific to whether the experience was intentional or not. We examined both evoked response that occur in response to events in the task, as well as oscillatory measures of neural activity that reflect patterns of ongoing neural processing.

## Materials and Procedure

### Subjects

Twenty-six participants (12 females, age M: 25 SD: 4.3) with no history of neurological or psychiatric disease volunteered or received partial course credits to participate in the study. All had normal to corrected-to-normal vision. All procedures were approved by Trinity College Dublin ethics committee and in accordance with the Declaration of Helsinki. Participants were informed extensively about the experiment and all gave written consent. They were seated in a sound-attenuated, electrically shielded and dimly lit room, at a viewing distance of approximately 70 cm of a 20” CRT monitor with a refresh rate of 60Hz.

### Fixed SART

During the fixed SART, digits from 1 to 9 were displayed in sequence and individually for 250 msec on the center of the monitor with a fixed inter-stimulus interval (ISI) of 2315 msec. The choice in ISI was the result of piloting with varying stimulus presentation rates prior to the experiment. The ISI used here represents the best choice to induce a maximum number of MW episodes^26^ without imposing excessive demands on endogenous attention by being too monotonous and exacerbating the task’s difficulty^89^.

The “no-go” target to which the participants were instructed to withhold response was the number 6 (11% of the trials) leaving the numbers 1 to 5 and 7 to 9 as “go” stimulus to which they were instructed to respond as fast as possible while avoiding anticipation responses. Stimulus were presented centrally at a font size of 140 in Arial using the Presentation software package (Version 19.0; www.neurobs.com). Participants were instructed to lock their response to the offset of the stimulus, a response strategy that has been successfully applied to minimize both the inter-individual variability in response times and the speed-accuracy tradeoffs^44,90^.

The duration of each of the three SART blocks lasted for a minimum of 8 and a maximum of 15 minutes for a total running time ranging from 24 min to 45 min with approximately 800 to 1200 presented SART stimuli respectively. All three blocks were interspersed by two 2 min long eyes-closed sessions during which participants were instructed to close their eyes and relax.

### Thought probes

Throughout the SART, intermittent and self-caught thought probes were used to assess patterns of ongoing thought. Probe-caught (PC) sampling prompts pseudo-randomly interrupted the task after 12, 18, 24, 30, 36, 42 or 48 trials (approx. 28, 42, 56, 76, 83, 97, 111 sec). PC probes consists of interruptions of the experimental task at random intervals prompting participants to reveal the content of their thoughts just prior to the interruption^91^. Participants were also instructed to interrupt the task by pressing SPACE if they became aware of a shift of attention away from the task. SC probes require participants to monitor their attention and to interrupt the task as soon as they become aware of their mind straying^92^. Once triggered, thought-probes presented participants with the question “Where was your mind just now?” and prompted them to classify their immediately preceding thoughts by choosing between five possible answers: “(1) on the task; (2) about the task (3) distracted by sensations (4) on future plans or past memories (dTUT) (5) daydreaming (sTUT)”. The correct assignment of mental activity to one of the five categories was assured by giving participants an extensive description and real-world examples for each category and ask them to relate them with their own experiences. Subsequently, they completed a short questionnaire with ten examples of mental content or situations to be classified in one of the five thought probe categories.

The duration of the SART blocks varied depending on each participant’s attentional capacity and responses to the thought probes. In the absence of SC probes, PC interruptions would occur on average every 76 sec resulting in approximately 6 to 12 probes per block (for 8 and 15 min respectively). To balance the distribution of off-task reports and obtain at least 8 reports per subjects for each of the relevant conditions to be compared during the analysis (on-task, dTUT and sTUT) each block would terminate at the earliest after 8 min only if 3 reports were recorded for the aforementioned conditions. Each block would terminate at the latest after 15 min regardless of the reports collected. Therefore, SART blocks lasted for a minimum of 8 and a maximum of 15 minutes for a total running time ranging from 24 min to 45 min with approximately 800 to 1200 presented SART stimuli respectively. All three blocks were interspersed by two 2 min long eyes-closed sessions during which participants were instructed to close their eyes and relax. The task ran for an average of 38.7 min (SD: 4.8) with an average of 65.3 thought probes (SD: 31.2) of which 71.4% were of the SC variant (for an overview see Figure 1).

**Figure 1.**
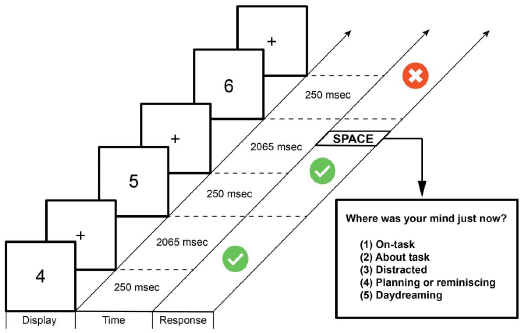
Fixed SART with thought probes. Participants were instructed to respond to each sequentially presented digit except for the target digit “6” to which they should withhold response. In addition, they were told to monitor their thought process and interrupt the task if they found themselves MW by pressing the SPACE key which triggered SC probe. PC probes would interrupt the task after a pseudo-random number of trials (12,18,24, 30, 36, 42 or 48). Both PC and SC probes prompted participants to categorize their attentional state.

### Data acquisition

64-channel EEG data was acquired at 512 Hz using a BioSemi ActiveTwo system (biosemi.com) placed according to the international 10-20 system. To assess eye movements and blinks, 4 electro-oculographic channels (EOGs) were placed above and beneath the left eye, and on the side of the left and right eye near the temples. The behavioral responses were recorded with the presentation software and markers were written unto the EEG recording with the help of the Biosemi system.

### Data processing

All data were processed, analyzed and visualized through Matlab (The Mathworks) with the help of custom written scripts and the following toolboxes: EEGLAB^93^, Fieldtrip^94^ and the BBCI toolbox (www.github.com/bbci/bbci_public).

### EEG data preprocessing

Offline EEG data was temporarily high pass filtered at 1Hz before line noise removal (50Hz) and robust referencing were applied with the PREP pipeline^95^. To avoid contamination of the data by noisy channels, the robust referencing algorithm identified, removed and interpolated those channels before referencing. The data was subsequently high-pass filtered at 0.5 Hz and downsampled to 250 Hz. Channels exhibiting activity above 200 standard deviations from the mean and previously interpolated channels were removed to prevent the introduction of nonlinearities for the subsequent independent component analysis (ICA). Sections of the data showing activity above 300 μV were trimmed. ICA, a blind-source separation method, was applied with EEGLAB 13 to extract independent components, which were subsequently inspected and automatically label with MARA^96^ and subsequently used to automatically identify and reject artifactual component (threshold for rejection was set at a conservative 95% probability to favor preservation of neural activity). Lastly, the data was visually inspected to remove remaining artefactual sections.

### EEG data segmentation

To categorize the SART stimuli according to the subject’s attentional state the EEG time-course was segmented into stimulus-locked 2.8 sec epochs (−200 to 2400 msec relative to SART stimulus) extending backwards four trials from the thought probe prompt. Although the temporal dynamics and the exact onset of attentional shifts remain unknown and difficult to assess^31^, we chose a window of four SART trials prior to a thought probe as reflecting the reported attentional state. This window, spanning a period of approximately 9.3 sec was chosen *a priori* for the following reasons:

1. a 10 sec time window has been used extensively in the prior analysis investigating MW^36,37,45,49^
2. Behavioral measures of MW, such as faster response times and increased variability, have been observed during the four SART trials leading to a commission error^7,86,97,98^
3. A 9.3 sec time-window would capture the entire duration of a specific attentional state, which has been suggested to last around 2.5 to 10 sec^99^, without extending too far back as to capture a prior state.

Each group of 4 epochs were subsequently associated with the attentional state according to the thought probe report. The aim of the study being the investigation of the neurophysiological difference between dTUT and sTUT, the TRI and distraction conditions were included in the design of the experiment to guarantee the accurate classification of different types of thought but were not included in the analysis. Due to the nature of the task, participants were only instructed to trigger a SC probe when they caught themselves off-task and as a result the SC condition lacked on-task reports.

The conditions were separated in the following manner. On-task condition was only considered in the context of PC reports (PC_OT). Due to the fewer number of trials for PC_dTUT and PC_sTUT, these two conditions were concatenated (PC_dTUT+PC_sTUT=PC_MW) as a general measure of off-task activity for comparison with PC_OT. Two SC conditions were considered, SC_dTUT and SC_sTUT.

In summary, analyses were run on concatenated 4 pre-trials for the following conditions pairs: (1) PC_OT/MW: probe-caught on-task (PC_OT) vs. probe-caught MW (PC_MW =PC_dTUT+PC_sTUT) and (2) SC_dTUT/sTUT: self-caught deliberate MW (SC_dTUT) vs. self-caught spontaneous MW (SC_sTUT). The exact distribution for the condition pairs can be taken from Table 1.

**Table 1.** Total number of thought probes and average per subject for each condition included in the analysis. For each thought probe

|  | PC_OT | PC_MW | SC_dTUT | SC_sTUT |
| --- | --- | --- | --- | --- |
| Total Nb | 95 | 84 | 313 | 300 |
| Average per Subject | 4.25 | 4.2 | 15.7 | 15 |

All trials that weren’t rejected due to artifacts were included in the average for the ERP analysis resulting in a total of 19 and 18 subjects for the condition pairs PC_OT/MW and SC_dTUT/sTUT, respectively. For subsequent analyses participants that did not have at least 8 trials (amounting to a total of 32 epochs) for both conditions of a condition pair were not considered which reduced the number of participants for the PC_OT/MW pair to 15.

### ERP analysis

Artifact-free epochs from −200 to 2400 msec with respect to the four SART stimuli prior to a thought sampling prompt (PC_OT/MW and SC_dTUT/sTUT) were extracted and averaged. A −200 to 0 msec prestimulus baseline was used for all ERP components analysis and visualisation. ERP component structures were confirmed by visual inspection of grand-average waveforms. The width of the temporal interval used to measure average component amplitudes were based on the spatial extent and approximate duration of each component. Visual inspection of the grand averaged waveforms revealed a tri-phasic complex (P1-N1-P2; 80-110, 120-170 and 200-300 msec respectively) over parieto-occipital (POcc) regions and a positive deflection over fronto-central scalp regions peaking around 120-170 msec (P1a) (see Figure 2 and 3). Parieto-occipital ERP contrasts were evaluated with 2*2*3 repeated-measures analysis of variance (ANOVA) with factors condition (COND) x channel location (LOC) x ERP component/ time interval (TIME). For simple contrasts, e.g., channel FCz with two conditions, and post-hoc testing, paired sample t-tests were conducted. The overall alpha level was set at 0.05. In the interest of brevity and relevance to the investigative aims of the study, only significant effects or differences are reported. To precisely determine channels and time-intervals exhibiting significant differences across conditions cluster-based permutation testing (allowing for a more accurate approximation of the Monte-Carlo statistic and correcting for multiple comparisons) was conducted as proposed by Maris & Oostenveld^100^ and implemented in Fieldtrip.

**Figure 2.**
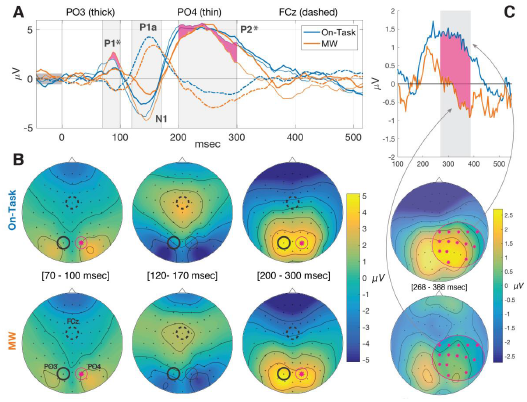
(A) Grand-average ERPs for condition-pair PC_OT/MW (collapsed 4 trials prior to PC interruption) for channels FCz, PO3 and PO4 with (B) scalp topographies for the marked intervals 80 to 100 msec, 130 to 260 msec and 200 to 300 msec after SART stimulus. (C) Averaged waveforms of the right centro-parietal channels identified during cluster-based permutation testing with corresponding scalp topographies for time interval 268 to 388 msec (pink shaded area and channels are statistically divergent p<0.05).

**Figure 3.**
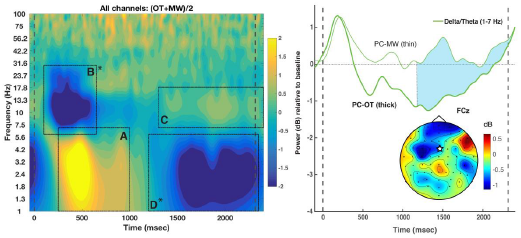
Left panel: Time-frequency representation (OT+MW)/2 for collapsed 4 trials prior to probe-caught thought-probe averaged over all non-boundary channels. The rectangles A to D correspond to the four global TFWs of interest containing the individually selected subject-specific windows. TFWs marked with an asterisk were found to differ significantly across conditions. Right panel: Time course of extracted delta/theta power relative to baseline for channel FCz for on-task (thick) and mind-wandering (thin). The blue shaded area corresponds to the window D which was found to be statistically divergent for channel FCz. Also shown for that time window is a topographical difference map for the delta/theta band activity (1 to 7 Hz) between on-task and mind-wandering (OT – MW).

First, t-values were subjected to a cluster-level test on the entire time course of the trials for the grand average ERPs of subjects for both conditions of a condition pair. The significance of the difference between two compared condition, the Monte-Carlo p-value, is estimated by comparing the observed cluster-level statistic with a randomization null distribution, obtained here from 3000 random draws, under the assumption of no differences between conditions. Finally, the significantly differentiating (p<0.05) cluster of channels and timepoints were subjected to t-tests and averaged for visualization.

### EEG time-frequency decomposition

The epoched EEG of the 4 collapsed pre-trials for both condition pairs PC_OT/MW and SC_dTUT/sTUT were baseline normalised against the 5th trial and subsequently decomposed in their time-frequency representation with custom written Matlab routines^101^. Complex Morlet wavelets, defined as Gaussian-windowed complex sine wave *e^i2πtf^e*^−*t*^2^^/(2*σ*^2^), where *t* is time, *f* is frequency (1 to 100 Hz in 100 logarithmically spaced steps), *σ* is the complex operator and *σ* defines the width (or the number of “cycles”) of each wavelet. *σ* set according to 2*πf*, increased linearly from 1 cycle for 1 Hz to 10 cycles for 100 Hz thus optimizing the tradeoff between temporal and frequency smoothing^102,103^. From the resulting analytical signal we extracted an estimate of frequency-band specific power and phase by squaring the magnitude of the convolution result *Z_t_* (*real*[*z_t_*]^2^ + *imag*[*z_t_*]^2^) and computing *ϕ_t_* = *arctan*(*imag*[*z_t_*]/*real*[*z_t_*]) respectively. For the decibel normalisation of power (*dB power* = 10 * *log*10[*power*/*baseline*]) we chose the trial immediately preceding the 4 collapsed trials of the condition pairs (the fifth trial counting back from the thought probe onset) as baseline. The phase or inter-trial phase coherence (ITPC) is a measure of the similarity in phase angle of the oscillation across trials hence bound between 0, no phase clustering, and 1, complete phase clustering across trials. The phase coherence value is defined as follows:

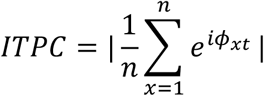

The statistical analysis of the average activity (time-frequency decomposition and ITPC) for each subject followed a selection procedure orthogonal to potential condition. To reduce the number of tests time-frequency representation were averaged for all channels save boundary channels (channels in close proximity to the face were excluded on the grounds of a higher susceptibility to artifacts). We inspected the time-frequency results of (COND A + COND B)/2 relative to baseline (fifth pre-trial) for each subject to determine time-frequency windows (TFWs) for comparison while avoiding circular inference. Visual inspection of the changes relative to baseline of the decomposed signal for all conditions revealed similar patterns of activity, two stimulus evoked changes: (A) an event-related desynchronization (ERD) in the alpha/beta band and (B) an event-related synchronization (ERS) in the delta/theta band, and two tonic changes: (C) an increase in activity centered on the alpha band and (D) a suppression in the delta/theta band. For each of these changes a global TFW was defined to account for slight variability in time and frequency distribution across condition-pairs. For the PC_OT/MW condition-pair we set the global TFW to span from 1 to 7 Hz in frequency and from 250 to 1000 msec relative to stimulus onset (Figure 3, A) for the delta/theta band ERS, 6 to 30 Hz and 100 to 700 msec for the alpha/beta band ERD (Figure 3, B), 7 to 18 Hz and 1300 to 2400 msec for the alpha increase (Figure 3, C), and lastly 1 to 8 Hz and 1200 to 2350 msec for the delta/theta suppression (Figure 3, D). For the SC_dTUT/sTUT condition-pair, we set the global TFW to span from 2 to 8 Hz in frequency and from 0 to 1000 msec relative to stimulus onset for the delta/theta band ERS (Figure 5, A), 6 to 40 Hz and 100 to 800 msec for the alpha/beta band ERD (Figure 5, B), 7 to 17 Hz and 700 to 2350 msec for the alpha increase (Figure 3, C), and 1 to 8 Hz and 1000 to 2350 msec for the delta/theta suppression (Figure 3, D). We then selected subject specific TFWs within each global TFW to account for inter-individual variability in peak frequencies of bands of interest^104–106^.

For each selected window all time-frequency points were averaged and submitted to a repeated-measure ANOVA (COND*TFW). We report Greenhouse-corrected p-values whenever sphericity was violated. Post-hoc two-tailed t tests were used to examine directional effects. To confirm results, we ran the same analysis on a priori chosen channels FCz and PO3/4, which are routinely selected to investigate FMT and alpha band activity respectively^107–109^.

### Data availability

The datasets generated and/or analyzed during the current study are available from the corresponding author on reasonable request.

## Results

### Probe-caught on-task vs. off-task

#### Event-related potentials (PCOT/TUT)

The CONDxLOCxTIME ANOVA for the pair PC_OT/MW revealed a significant main effect for the LOC [F(1,18)=5.32, p=.033]and TIME [F(2,36)=22.33, p<.000], as well as significant interaction effects CONDxLOC [F(1,18)=11.42, p=.004] and LOCxTIME nearing significance [F(2,36)=3.01, p=.058] (Greenhouse corrected p=0.07). Post-hoc comparisons showed decreased amplitude for the MW condition on right Parieto-occipital channel PO4, with P1 (OT M:1.97μV SD: 3.57; MW M: 0.57μV SD: 3.28)[t(18)=2.348 p=.03] and P2 (OT M:4.52μV SD: 4.24; MW M: 3.02μV SD: 4.75)[t(18)=2.38 p=.028] reaching and N1 (OT M:-2.53μV SD: 3.53; MW M: −3.83μV SD: 3.78)[t(18)=1.97 p=065] approaching significance. A t-test was conducted on the cluster of channel identified during the Monte-Carlo permutation (Pz, CPz, Cz, C2, C4, C6, T8, TP8, CP6, CP4, CP2, P2, P8 and PO4) for the time-interval 268 to 388 msec revealing a significantly decreased activity during the MW condition over right Parieto-occipital areas (OT M:1.32μV SD: 1.41; MW M: −0.28μV SD: 2.03)[t(17)=4.33 p<.001] (see Figure 2 panel C).

To sum up, the PC_OT/MW condition pair revealed a right parieto-central attenuation as observed by a reduction in P1 and P2 amplitude over PO4 and reduced amplitude over parieto-central sites for the P3 time-interval (268 – 388 msec; see Figure 2).

#### Time-frequency analysis (PC_OT/MW)

The 4×2 Time-Frequency Window x Conditions (TFWxCOND) ANOVA on all averaged channels returned a significant main effect for TFW [F(3,42)=42.39, p<.000] and a significant main effect for COND [F(3,51)=3.46, p<.013]. T-tests unveiled a significantly decreased alpha-beta ERD (Figure 3, B) for MW pre-trials (M:-2.27 dB SD: 1.55) in comparison with OT pre-trials (M:-1.13 dB SD: 1.26) [t(14)=2.27 p=.039]. This difference was exacerbated for PO3/4, [t(14)=2.94 p=.011], MW pre-trials (M:-3.49 dB SD: 1.72) vs. OT pre-trials (M:-1.94 dB SD: 1.25). Delta/theta power (thereafter referred to as FMT) measured at FCz [t(14)=2.45 p=.028], showed a significant decrease for the TUT condition (M:-2.69 dB SD: 1.24) relative to OT (M:-1.89 dB SD: .95) (Figure 3, right panel). The difference in tonic alpha power did not reach statistical significance for any of the tested channel constellations.

T-tests for the ITPC values for all three sets of electrodes tested (all averaged non-boundary channels, FCz and PO3/4) failed to return significant differences for the PC_OT/MW condition pair.

In summary, the on-task vs. off-task (MW) comparison revealed a right-lateralized attenuation for the components P1 and P2 ERPs over channel PO4, and for the P3 time-interval (268 – 388 msec) over parieto-occipital sites (Figure 2). Additionally a significant increase in power relative to baseline was found for the ERD (Figure 3, B), tonic theta power (Figure 3, D) and FMT (Figure 3, right panel) during off-task.

### Deliberate vs. spontaneous mind-wandering

#### Event-related potentials (SC_dTUT/sTUT)

The CONDxLOCxTIME ANOVA returned a significant main effect on TIME [F(2,34)=10.95, p<.000] and significant interaction effects CONDxLOC [F(1,17)=7.3, p=.015], CONDxTIME [F(2,34)=22.06, p<.000], LOCxTIME [F(2,34)=24.78, p<.000]and CONDxLOCxTIME [F(2,34)=21.17, p<.000]. Post-hoc t-tests showed significantly decreased amplitude for the sTUT condition for the P2 component on both PO3 (dTUT M:5.01μV SD: 4.41; MW M: 3.64μV SD: 3.01)[t(17)=2.4 p=028] and PO4 (dTUT M: 4.44μV SD: 3.73; MW M: 3.56μV SD: 2.62)[t(17)=2.61 p=.018]. P1 amplitude difference on channel PO4 approached statistical significance (dTUT M: 1.79μV SD: 2.41; MW M: 0.75μV SD: 1.82)[t(17)=1.96 p=066].

The t-tests conducted on the averaged amplitudes of the two sets of channels the over the time-interval 178-304 msec (fronto-central: AF7, AF3, F1, F3, FC3, FC1, Fpz, Fp2, AF8, AF4, AFz, Fz, F2-8, FC6, FC4, FC2, FCz; parieto-occipital: P1-9, PO7, PO3, O1, Oz, POz, Pz, P10, PO8, PO4, O2) identified with cluster-based permutation testing on all channels, revealed a significantly decreased amplitude for SC_sTUT over both fronto-central channels (dTUT M: −2.67μV SD: 2.53; sTUT M: −1.51μV SD: 2.08) [t(17)=-2.72 p=014] and parieto-occipital channels (dTUT M: 3.21μV SD: 3.15; sTUT M: 1.85μV SD: 2.85) [t(17)=3.28 p=005] (see Figure 4 panel C).

**Figure 4.**
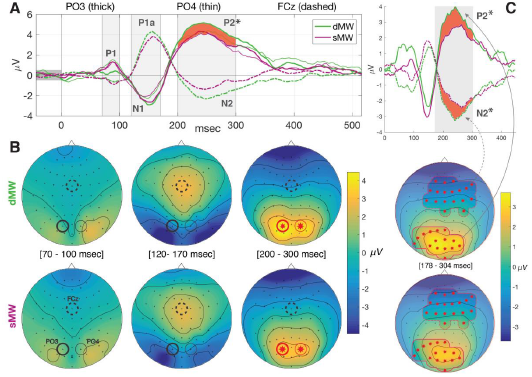
(A) Grand-average ERPs for condition-pair SC_dTUT/sTUT (collapsed 4 trials prior to SC interruption) for channels FCz, PO3 and PO4 with (B) scalp topographies for the marked intervals 80 to 100 msec, 130 to 260 msec and 200 to 300 msec after SART stimulus. (C) Averaged waveforms of the fronto-central and parieto-occipital channels identified with cluster-based permutation testing, with corresponding scalp topographies for time interval 178 to 308 msec (reddish shaded areas and channels are statistically divergent p<0.05).

To sum up, the P2 and N2 amplitudes were found to be significantly increased during dTUT.

#### Time-frequency analysis (SCdTUT/sTUT)

The 4×2 (TFWxCOND) ANOVA returned a significant main effect for TFW [F(3,51)=35.52, p<.000] and a significant interaction effect [F(3,51)=3.46, *p*<.023] (Greenhouse corrected p=0.46). Follow-up t-tests determined alpha activity (Figure 5, C and supplemental Figure 2, right panel) to be significantly higher during dTUT pre-trials (M:1.54 dB SD: 1.44) when compared to sTUT (M:.58 dB SD: 1.2) [t(17)=2.25 p=.038]. This contrast was slightly more pronounced when testing on FCz alone [t(17)=2.38 p=.029] with increased activity for dTUT (M: 1.46 dB SD: 1.1) relative to sTUT (M: .58 dB SD: 1.23) and remained significant for PO3/4 [t(17)=2.12 p=043] (dTUT, M: 2.43 dB SD: 2.07; sTUT, M: 1.08 dB SD: 1.55).

**Figure 5.**
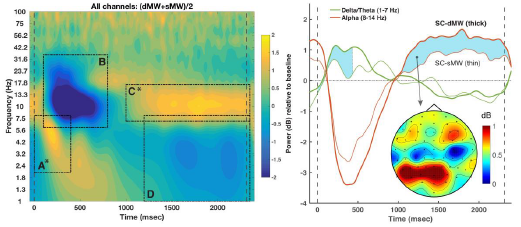
Left panel: Time-frequency representation (dTUT+sTUT)/2 for collapsed 4 trials prior to self-caught though-probe averaged over all non-boundary channels. The rectangles A to D correspond to the four global TFWs of interest containing the individually selected subject-specific windows. TFWs marked with an asterisk were found to differ significantly across conditions. Right panel: Time course of extracted alpha and delta/theta power relative to baseline for all non-boundary channels for deliberate (thick) and spontaneous mind-wandering (thin). Differences that were found to be differ statistically are shaded in blue. Also shown is a topographical difference map for the alpha band activity (8 to 14 Hz) between dTUT and sTUT (dTUT-sTUT).

The t-test for the delta-theta ERS on all channels (Figure 5, A) approached statistical significance [t(17)=2 p=.062], hinting towards an increase for dTUT (M: 1.53 dB SD: 1.1) relative to sTUT (M: .92 dB SD: 1.25). The difference reached statistical significance for PO3/4 [t(17)=2.2 p=042] with dTUT (M: 2.35 dB SD: 1.5) and sTUT (M: 1.56 dB SD: 1.42). However, theta activity (Figure 5, D) did not exhibit statistically significant differences for any of the three channel configurations.

Lastly, two-tailed t-tests were found to be significant for ITPC values for all averaged channels [t(17)=2.8 p=.012] with increased coherence for dTUT (M: .52; SD: .1) relative to sTUT (M: .45; SD: .1), and PO3/4 [t(17)=2.94 p=.009] also with increased coherence for dTUT (M: .64; SD: .15) and sTUT (M: .58; SD: .16) with FCz nearing significance [t(17)=2.09 p=.052] (see supplemental Figure 3).

In summary, dTUT exhibited significantly more extensive amplitudes for the P2 time interval over both central and parieto-occipital sites, and channels PO3/4. Crucially, we found increased tonic alpha power (and ERS) for dTUT compared to sTUT (see Figure 5). A complete summary across the two condition pairs compared can be taken from Table 2.

**Table 2.**
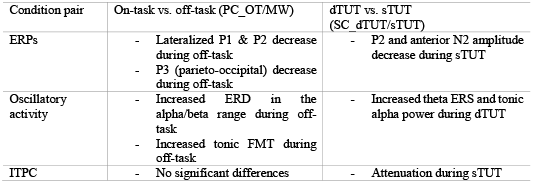
Overview of the significant differences found for the comparison of the condition pairs PC_OT/MW (on-task vs. off-task) and SC_dTUT/sTUT (deliberate vs. spontaneous TUT)

## General Discussion

Our study provides neurocognitive evidence that off-task states with different experiential features depend on a different balance of underlying neurocognitive processes. Replicating prior studies, we found that periods of off-task thought reported at probes were associated with reduced attention to external task relevant information. In particular we found reductions in the amplitude of aspects of the event related potential approximately 100-300ms post stimulus onset during the off-task state. These results support prior studies that show reduced processing of stimuli in this time range for both online and offline measures of task unrelated thinking^26,36,38,39^. Together these results are consistent with the hypothesis that reduced processing of external task relevant information occurs whenever attention shifts from the task in hand. Importantly, we also found evidence that patterns of neural activity distinguish whether off-task thought was experienced as deliberate or spontaneous. In particular, we found that greater power in alpha and theta bands in the periods when off-task was described as deliberate. In contrast spontaneous periods of off-task thought were linked to more pronounced reductions within early evoked components occurring around 200 milliseconds of stimulus onset. Consistent with component process accounts of ongoing thought these data suggest that periods of off-task thought with different experiential features depend on a different balance of neural processes. We consider the implication of these results for contemporary accounts of ongoing thought.

Our data suggests that low frequency oscillations within the EEG are particularly prevalent during periods when off-task thought is engaged in a deliberate manner. Oscillatory activity within the alpha range is thought to be important for attentionally demanding tasks, and particularly those that rely heavily on internal information^110^. Importantly, alpha activity has been shown to be important in tasks of creativity^111^, a well-documented correlate of trait variation in off-task states^25,112,113^. One hypothesis is that alpha activity elicits its effects on cognition role through a top down process that organizes neural function by inhibiting irrelevant sensory input^114^.

In contrast, spontaneous episodes of off-task thought were associated with a more pronounced reduction in evoked responses, as well as less coherence in brain activity across time. Periods of spontaneous off-task thought were associated with reductions in positive amplitude evoked responses focused on the occipital cortex (the P2), as well as reductions in negative amplitude components with a fronto-central locus (the N2). The amplitude of the P2 over the occipital cortex is generally assumed to reflect higher order perceptual processing of visual stimuli^115^. In addition, periods of spontaneous off-task thought were linked to reductions in the amplitude of the N2, a component that is thought to reflect processes linked to executive control such as conflict monitoring^116^. The combination of reduced P2 and N2 linked with spontaneous off-task thought suggests that this aspect of off-task experience may be associated with failure in the top down maintenance of the appropriate task set. This interpretation seems to be consistent with prior studies that suggest that spontaneous off-task thought is primarily linked to lack of external alertness and lower executive control^117^.

More generally, our findings suggest that the degree of intentionality during off-task thought can be understood as the interplay between top-down and bottom-up features. We found that when participants were engaged in deliberate off-task thought, alpha activity was increased, a pattern that is often observed when individuals are deliberately engaged in internal tasks^71,110,118^, and may reflect patterns of top down inhibition of irrelevant sensory input. In contrast, periods of spontaneous off-task thought were linked to evoked response that may indicate the higher order processing of sensory input and conflict monitoring necessary to detect failures in task set. Together these patterns are broadly consistent with the hypothesis that spontaneous and deliberate off-task thought reflect differences in the role that top down processes play in ongoing thought. In particular, based on our data we may expect that deliberate off-task thought is linked to the motivated maintenance of a task set, while more spontaneous off-task episodes are linked to intermittent failures to maintain task set in attention. The role of executive control in off-task thought has been a central question in the literature, with evidence for a role in both task-related and off-task thought. In this context, our neural data suggests that the level of intention in the off-task state may be a key moderating factor that determines whether the experience is linked to a failure in control or relies on the same resource to organize an internal train of thought.

## Acknowledgments

This publication was supported by a Marie Curie Initial Training Network grant (n° 606901) under the European Union’s Seventh Framework Programme. JS was supported by the European Research Council (WANDERINGMINDS 646-927). PD was supported by the Irish Research Council (IRC Laureate grant: IRCLA/2017/306).

## Authors contribution

A.M., P.D. and I.R. designed the study, A.M. prepared the experimental paradigm and collected the data, A.M., M.A. and P.D. participated in data analysis, A.M. drafted and J.S. revised the manuscript. All authors discussed the results and commented on the manuscript.

## Competing Interests

The authors declare no competing interests.

---

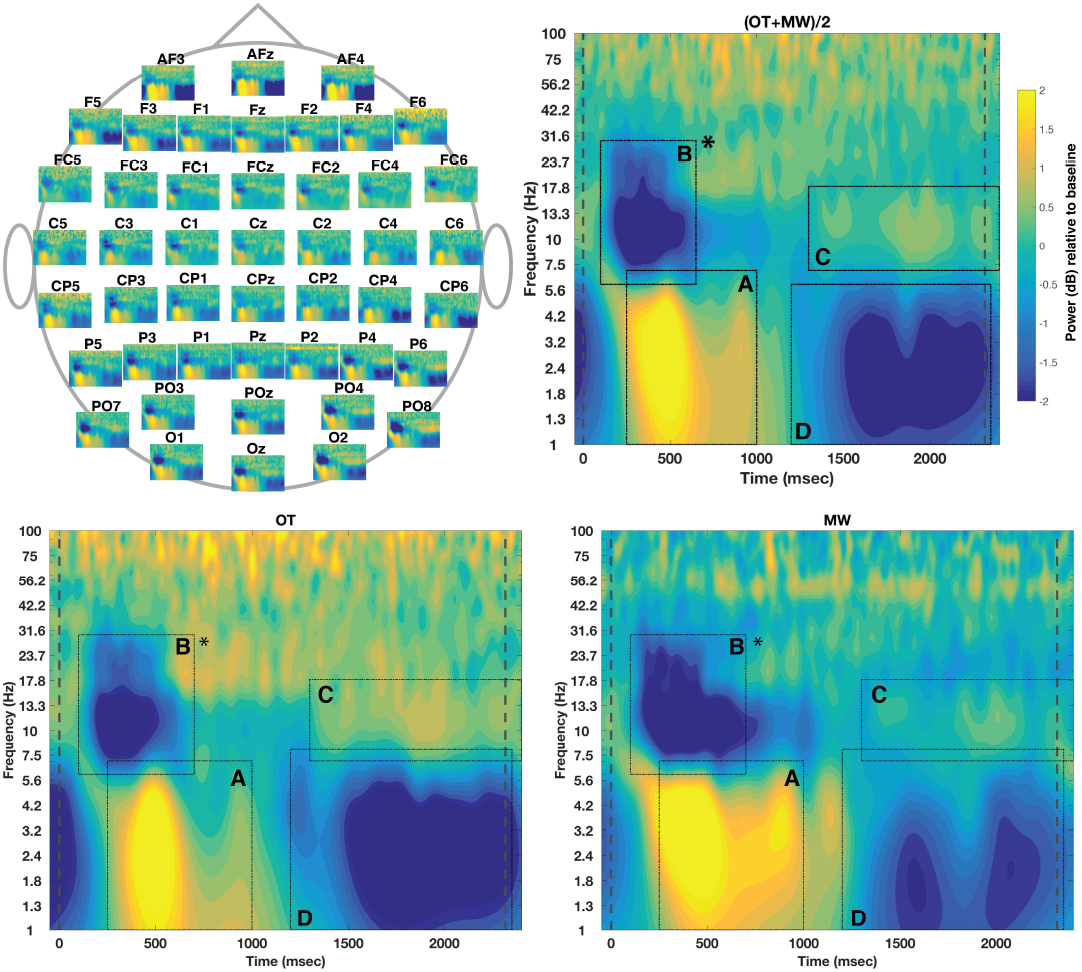

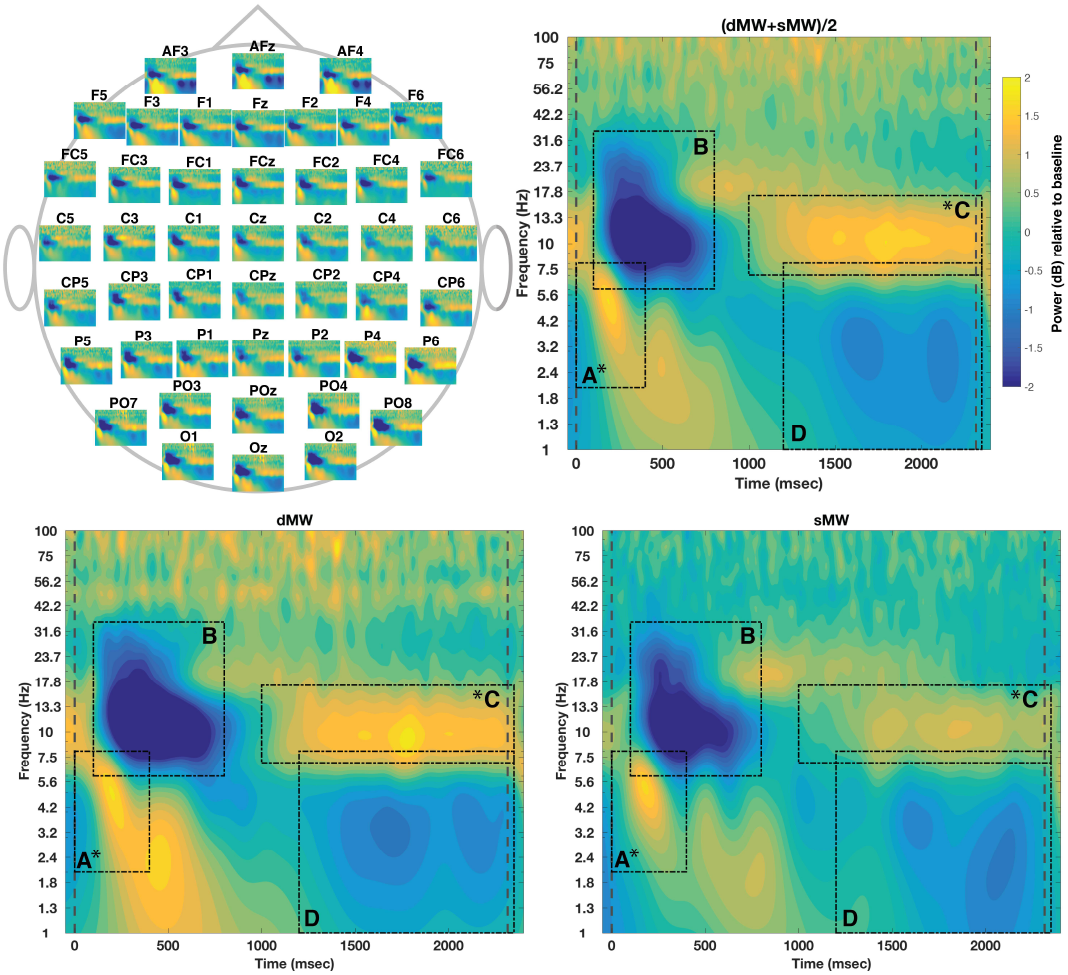

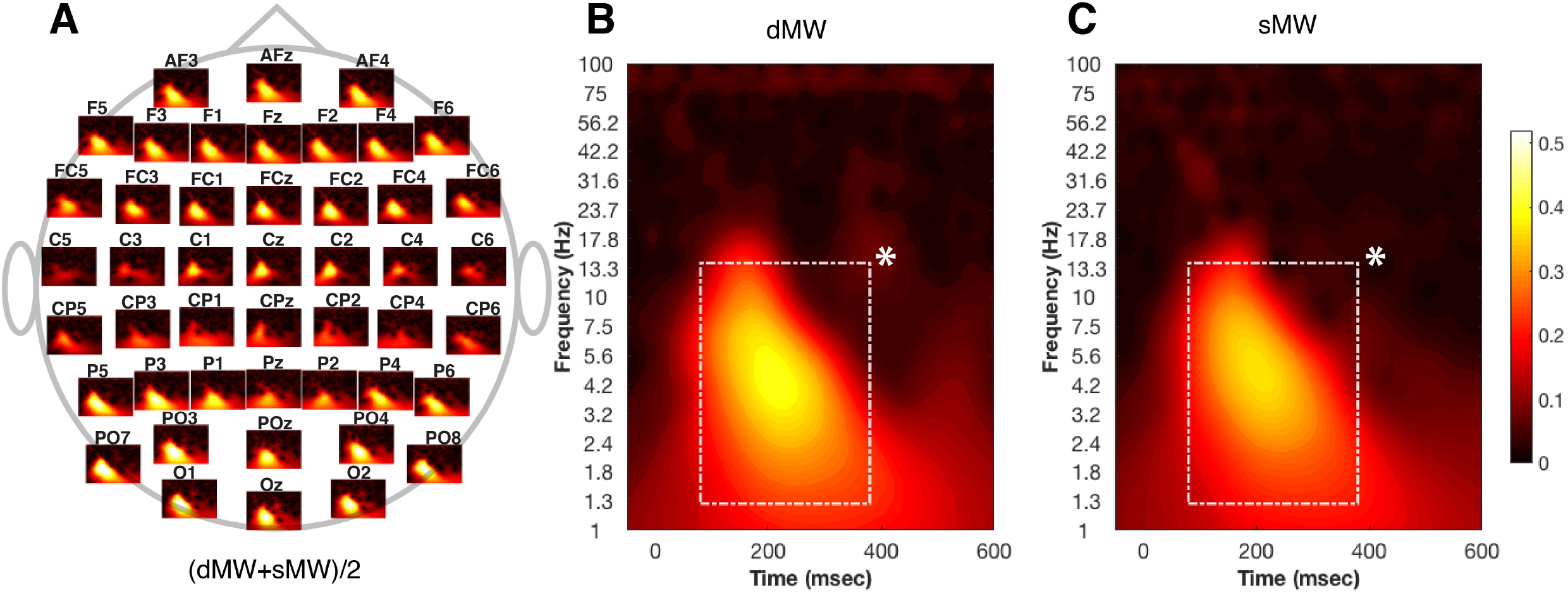

